# South African brown locusts, *Locustana pardalina*, hosts fluconazole resistant *Candidozyma* (*Candida*) *auris* (Clade III)

**DOI:** 10.1101/2025.01.22.634428

**Authors:** Adepemi Ogundeji, Maryam Bello-Akinosho, Vaughn Swart, Jonathan Featherston, Armand Bolsenbroek, Carel Beneke, Jolly Musoke, Tyla Baker, Arshad Ismail, Olihile Sebolai, Jacobus Albertyn, Carolina Pohl

## Abstract

The environmental niche and mode of transmission from the environment to humans of the emerging pathogenic yeast, *Candidozyma* (*Candida*) *auris* is a subject of speculation, with hypotheses including avian species and marine environments. Interestingly, yeasts related to *C. auris* have been repeatedly observed associated with various insects. This lprompted us to investigate a thermophilic insect, *Locustana pardalina* as possible host for *C. auris*. Here we report the isolation and identification of three *C auris* strains from the gut of *L. pardalina* as well as the phenotypic characterisation of one of these isolates. Interestingly, the isolate was able to survive at 50oC and grew at 15% NaCl. In addition, it was susceptible to the tested disinfectants and antifungals, except fluconazole. Genome sequencing and SNP analyses placed the isolate in Clade III, which is common is South Africa. This highlights the role of insects in the evolution and dissemination of emerging pathogenic yeasts.

## Introduction

*Candidozyma* (*Candida*) *auris*, an emerging pathogenic yeast that is able to cause nosocomial outbreaks, forms part of a clade of closely related species, consisting of *C. auris, C. duobushaemulonii, C. haemulonii, C. haemuli var. vulneris, C. heveicola, C. khanbhai, C. konsanensis, C. metrosideri, C. ohialehuae, C. pseudohaemulonii, C. ruelliae* and *C. vulturna* in the *Metschnikowiaceae*. This classification has recently been revised, and these species have been reassigned to the new genus *Candidozyma*. Many of these species can cause human infections and are resistant to antifungal drugs (Liu et al., 2024).

One of the unanswered questions regarding the emergence of *C. auris* is its environmental niche and mode of transmission from the environment to humans. It has been hypothesised that this yeast might have intermediate hosts, such as birds, which could facilitate its spread and transmission to humans (Casadevall et al. 2019). This is speculated to be due to the high body temperature of birds that would allow infection only by thermophilic yeasts able to grow at >40oC, and to then be spread via migratory birds to different geographic locations, although no evidence is available regarding direct isolation of *C. auris* or the presence of its DNA from birds (Akinbobola et al. 2023, Irinyi et al. 2023).

Interestingly, members of *Candidozyma* do seem to be associated with insects (among other niches), which may act as vectors of these yeasts. Strains of *C. duobushaemulonii* have been isolated from the European firebug (*Pyrrhocoris apterus*) (Cendejas-Bueno et al., 2012), and its DNA detected in microbiomes from the Asian tiger mosquito (*Aedes albopictus*). Similarly, DNA sequences of *C. haemulonii* were present in Svensson’s copper underwing (*Amphipyra berbera*), the cosmopolitan springtail (*Entomobrya nivalis)*, double-spined bark beetle (*Ips duplicatus*), six-toothed bark beetle (*Ips sexdentatus*), slender springtail (*Orchesella flavescens*) and longhorn crazy ant (*Paratrechina longicornis*). In addition, sequences of *C. haemulonii, C. haemulonii var. vulneris and C. duobushaemulonii* were abundant in microbiomes of termites (*Isoptera* spp.), while DNA sequences of *C. helveicola* were detected in samples from the gut of the Julia butterfly (*Dryas iulia*), the ghost yellow butterfly (*Eurema albula*) and the spot-banded daggerwing (*Marpesia merops*) (Iriny et al. 2023). This insect association prompted us to investigate the presence of *C. auris* (a known thermotolerant yeast) in insects that can withstand high temperatures.

An example of a group of cosmopolitan insects with high body temperature preference is locusts. Migratory locusts (*Locusta migratoria*) have a preferred body temperature of 38 – 39°C if food is abundant (Miller et al., 2009), although lower temperatures are selected when food becomes limiting (Coggan et al., 2011). The desert locust (*Schistocerca gregaria*) can tolerate 50oC without apparent adverse effects (Maeno et al., 2023). Similarly, the brown locust (*Locustana pardalina*) is typically exposed to high ambient temperatures of 33 – 38°C and soil temperatures of 39 – 62°C and is reported to have a preferred body temperature of 39 –41oC (Duncan & Hanrahan, 2018). Thus, we investigated the presence of *C. auris* in adult *L. pardalina* in South Africa.

## Materials and methods

### Collection of *L. pardalina*

Twenty gregarious adult locusts were collected by sweep net on 16 April 2022 during a large locust outbreak, which occurred fromSeptember 2021 to May 2022. The coordinates of the sample site are 31°58’45.99”S and 24°42’7.58”E, within the semi-arid Eastern Karoo climactic region in the Eastern Cape of South Africa. This is a remote rural area, consisting mostly of large private grazing farms with minimal human occupation.

### Yeast isolation from the alimentary canal of brown locust

The locusts were surface sterilised, and the entire alimentary canal was removed and dissected into the fore-, mid-, and hindgut. Each section was vigorously rinsed in sterile distilled water, which was plated onto Yeast Malt Extract (YM) agar (3 g/L malt extract, 3 g/L yeast extract, 5 g/L peptone powder, 10 g/L D-glucose, 16 g/L agar) and incubated at 30ºC. Colonies with different characteristics were sub-cultured until pure cultures were obtained.

### Identification of isolated strains and phylogenetic analysis

Both the ITS region and the D1/D2 domains of the LSU rRNA gene were amplified by PCR using the respective primer pairs of ITS4 (5′-TCCTCCGCTTATTGATATGC-3′) and ITS5 (5’-GGAAGTAAAAGTCGTAACAAGG-3’) as well as NL1 (5′-GCATATCAATAAGCGGAGGAAAAG-3′) and NL4 (5′-GGTCCGTGTTTCAAGACGG-3′), according to Kurtzman and Robnett (1998). Amplicons obtained were sequenced using the BigDye™ Terminator Sequencing kit on the Applied Biosystems 3500 genetic analyser, following amplicon clean-up and post clean-up PCR. Consensus sequences obtained for each isolate were compared to the GenBank nucleotide data library using the Basic Local Alignment Search Tool (BLAST) software (Altschul et al., 1997) at the National Centre for Biotechnology Information (NCBI) website (http://www.ncbi.nlm.nih.gov).

All strains were deposited in the Biodiversity Biobanks South Africa (BBSA) Yeast culture collection, held at the Department of Microbiology and Biochemistry, University of the Free State and preserved in glycerol stocks at -80oC.

### Phenotypic characterisation Salt tolerance assay

*Candida auris* was inoculated into 5 mL Yeast Potato Dextrose (YPD) media (10 g/L yeast extract, 20 g/L peptone, 20g/L glucose) and incubated for 24h at 37°C while shaking. A 10x cell dilution was prepared with YPD media and the OD_600_ was measured with a Jenway 6400 spectrophotometer. Cells were standardised to an OD_600_ of 0.8. A 10X serial dilution series was prepared for the standardised cells and 10 µL of the 10^−1^ to 10^−4^ dilutions were spotted onto prewarmed YPD plates supplemented with NaCl concentrations of 10%, 15% and 20%. These plates were incubated at 37°C for 72h.

### Temperature tolerance determination

*Candida auris* was grown and standardised to an OD_600_ of 0.8 and 200 µL of the standardised cells were plated into a 96-well plate with YPD negative controls. Growth at 30°C, 40°C and 50°C was monitored for 24 h in a Victor Nivo plate reader. This was done in triplicate and each 96-well plate contained five technical repeats.

### Antifungal susceptibility testing

Antifungal susceptibility, determined by the minimum inhibitory concentration (MIC), was assessed using the AST-YS08 card on the Vitek 2 Compact system (Biomerieux, France), in accordance with the manufacturer’s guidelines. The antifungal panel included fluconazole, voriconazole, caspofungin, micafungin, amphotericin B, and flucytosine.

### Disinfectant susceptibility testing

*Candida auris* was grown in YPD at 37°C overnight while shaking and standardised to 10^6^ cells/mL. A 2x serial dilution of each disinfectant was prepared in 2x RPMI-1640 of in a 96 well plate. Cells were then exposed to the disinfectant for 20 minutes and reinoculated in 2x RPMI-1640 to determine MIC values. The disinfectants used were absolute ethanol, 10% povidine iodine, 3.5% sodium hypochlorite, 12% polydimethylammonium chloride, 80% didecyldimethylammonium chloride, 50% benzalkonium chloride, 5% chlorhexidine gluconate and 0.5% chlorhexidine gluconate combined with 70% isopropyl alcohol.

### Genomic characterisation DNA extraction

*C. auris* UOFS Y-2024 was cultured in YM broth at 37°C overnight while shaking. Two ml of the culture was concentrated using centrifugation (Centrifuge 5430R Eppendorf® USA: F – 35 – 6 – 30 rotor) and was resuspended in 200 µl of Phosphate Saline Buffer. The complete 200 µl of resuspended cells were used to extract genomic DNA (gDNA) using the protocol stipulated in the ZYMO research Quick DNA TM Fungal/Bacterial Miniprep Kit (Orange; California). gDNA extraction was confirmed using gel electrophoresis (0.8% agarose gel at 90V, 400 mA for 25 minutes).

### Library Preparation, and Sequencing

Extracted genomic DNA was quantified using the Qubit 4 Fluorometer with the Qubit™ dsDNA HS Assay Kit (Thermo Fisher Scientific, Waltham, MA, USA). Paired-end library (2 × 150 bp) was prepared using Illumina DNA Prep kit (Illumina, San Diego, US), followed by sequencing on an Element Biosciences Aviti™ Sequencer using Cloudbreak Freestyle chemistry kit (Element Biosciences, San Diego, CA, USA) with 100 x coverage.

### Bioinformatics Analysis

Raw sequencing reads were processed using the v1.5 mycoSNP (Bagal et al., 2022) Nextflow (v23.10.0) nf-core (Ewels et al., 2020) pipeline. The Centre for Disease Control (CDC) *C. auris* reference genome (GCA_016772135.1_ASM1677213v1_genomic.fna) was used for alignment and single-nucleotide polymorphism (SNP) identification. In addition to the study isolates, based on clade-level classification by Suphavilai and co-worker (2024) sequence reads were obtained from the NCBI Sequence Read Archive (SRA) for a total of 32 publicly available *C. auris* genomes from clades I to VI and included for phylogenetic analyses (Supplementary Table 1). Briefly, (i) the mycoSNP pipeline prepared the reference geneome for alignment with nucmer, samtools, Picard tools and BWA index, (ii) sample QC was assessed with FastQC (Andrews, 2010), poor quality sequence data were trimmed with FaQCS v 2.10 and SeqKit, trimmed sequence reads were aligned to the reference with BWA MEM (v0.7.17), alignment file filtering with Picard (http://broadinstitute.github.io/picard/). The GATK4 pipeline (v4.2.5.0) was used for variant calling and to generate a multi-fasta from a combined variant calling file (VCF) using Bcftools (v1.14) and finally phylogenetic trees were drawn using fasttree (v2.1.10), quicksnp (v1.0.1) (https://github.com/k-florek/QuickSNP), iqtree (v2.1.4), rapidnj (v2.3.2) (Kurtz et al., 2004, Li et al., 2009, Andrews., 2010, Lo et al., 2014, Shen et al., 2016 Van der Auwera and O’Connor, 2020, Danecek et al., 2021, Price et al., 2010, Nguyen et al., 2015, Simonsen et al., 2008).

**Table 1.**
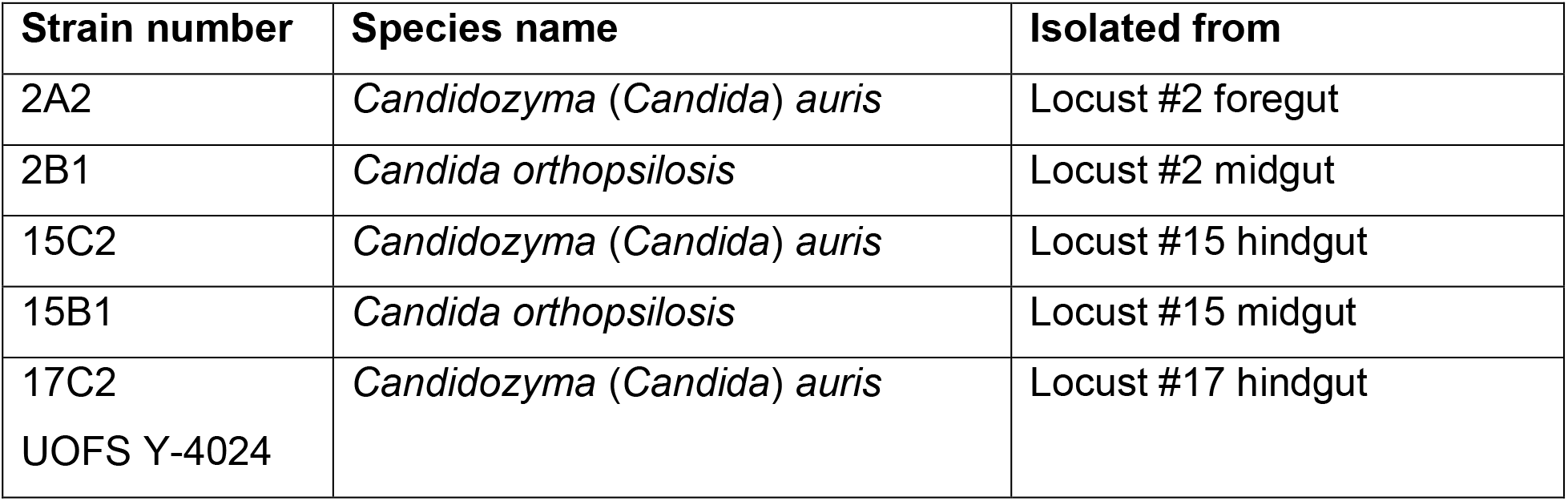
Potentially pathogenic yeasts isolated from the alimentary canal of *L. paradalina*.

## Results and Discussion

Three *C. auris* strains were isolated from three different adult locusts, two of which also harboured *Candida orthopsilosis* strains (Table 1). The fact that we were able to isolate *C. auris* from three locusts (15% of locusts) using non-selective media and a non-restrictive temperature of 30oC may indicate that *C. auris* is abundant in the locusts and that specific selective isolation is not mandatory.

Interestingly, although *C. orthopsilosis* was only isolated from the midgut, *C. auris* was isolated from the fore- and hindgut. Isolation from the foregut, which is dedicated to food intake and storage, filtering and partial digestion, indicates that *C. auris* was probably obtained by the locusts via feeding activities. Isolation from the hindgut confirms that *C. auris* can survive the digestive processes in the midgut and is likely to be released back into the environment via faeces (Stefanini, 2018). This and other studies (Dimitrov et al. 2024) highlight the importance of insects as potential vectors for the distribution of pathogenic yeasts.

We were able to revive one of the strains (17C2 = UOFS Y-4024) from cryostorage and phenotypic and genotypic analyses were performed on this strain.

### Osmo- and temperature tolerance of *C. auris* UOFS Y-4024

As expected, *C. auris* UOFS Y-4024 is salt tolerant and can grow on 15% NaCl (Figure 1). Interestingly, locusts (*Schistocerca* or *Locusta* spp.) require between 1 and 2% salts in their diet and can tolerate excess salts (up to 11%) and wide variations in the ratios of ions in the diet (Dadd, 1961). *L. pardalina* hoppers are also attracted to NaCl and NaH_2_PO_4_ which may be consumed, possibly to alleviate mineral deficiencies in their grass diet (Du Plessis & Botha, 1939). Although the soil and groundwater in the area where the locusts were collected is not particularly high in sodium salts and is considered slightly saline (200-400 mS/m) (Nell and Huyssteen, 2014; Nell, 2021), *L. pardalina* can travel up to 2.8 km per day (depending on the instar) while the flying adults can cross large distances (Duncan and Hanrahan, 2018). Thus, it is unknown how far they may have migrated during their swarming behaviour or where they may have ingested the yeasts isolated from them.

**Figure 1.**
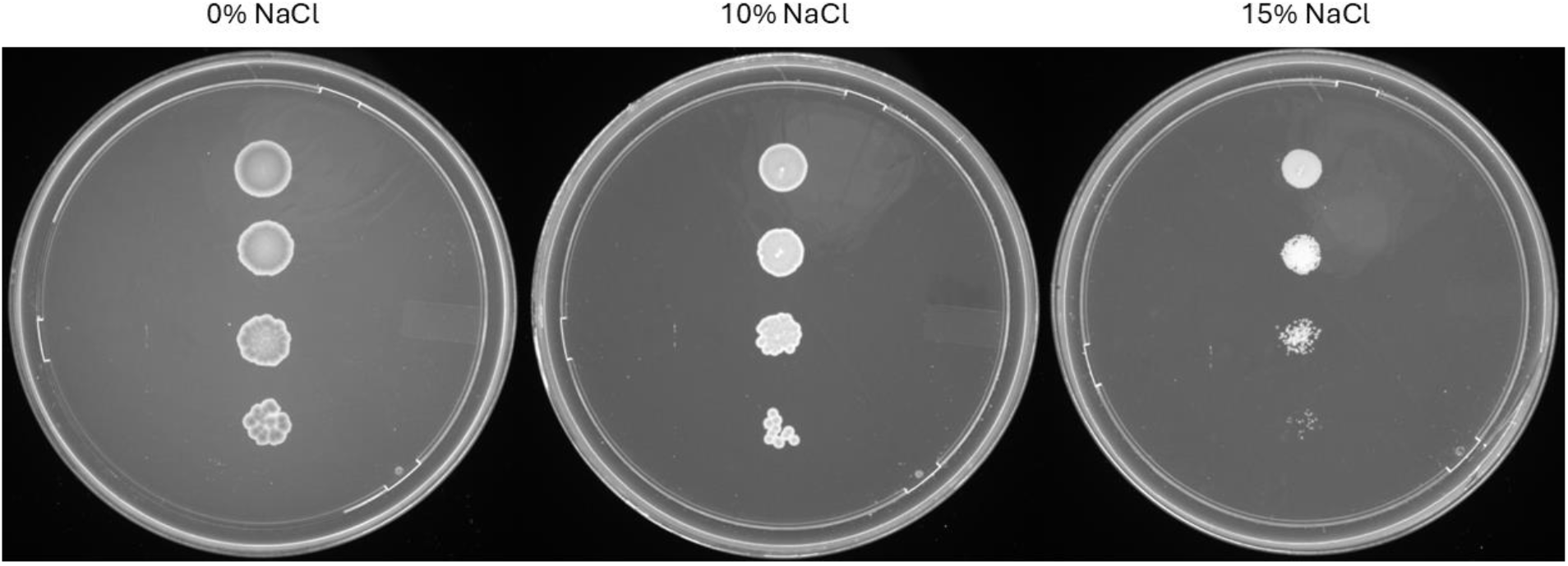
*C. auris* UOFS Y-4024 growth on NaCl concentrations ranging from 0% to 15 %

The isolate grew faster at 40^°^C, with a µ_max_ of 0.061h^−1^ after 120 min, compared to 30^°^C, where a lower µ_max_ of 0.044h^−1^ was obtained after 210 min. Interestingly, it survived and grew slowly (µ_max_ = 0.009h^−1^) at 50^°^C. This level of growth was attained after 210 min and maintained for an additional 240 min. This indicates that *C. auris* UOFS Y-4024 can grow well at the preferred body temperature of *L. pardalina* and can survive high soil temperatures typical of the Karoo region.

### Antifungal and disinfectant susceptibility of *C. auris* UOFS Y-4024

*C. auris* UOFS Y-4024 is resistant to fluconazole, with an MIC of 32 µg/mL as determined by Vitek 2 (Siopi et al., 2024), but susceptible to the other antifungals tested (Table 2). The decreased susceptibility to fluconazole is a common feature of many *C. auris* strains, including ones from non-clinical environments. For example, a *C. auris* strain isolated from an Egyptian cobra from a Moroccan market (also an arid environment) was also considered fluconazole resistant (Cafarchia et al., 2024).

**Table 2.** Minimum inhibitory concentrations (MICs) of antifungal drugs against *C. auris* UOFS Y-4024.

| Antifungal drug | MIC (µg/mL) |
| --- | --- |
| Fluconazole | 32 |
| Voriconazole | 1 |
| Caspofungin | 0.25 |
| Micafungin | ≤0.06 |
| Amphotericin B | ≤0.25 |
| Flucytosine | ≤1 |

The isolate was also evaluated against disinfectants from multiple classes (Table 3). Currently, no standards or recommended methods exist to eradicate *C. auris* from contaminated surfaces and infected individuals. Previously, ethanol has been shown to be effective against *C. auris* at a concentration of 70% (Rutala et al., 2019), and our results support this finding as the MIC determined was 29.5%. Similarly, povidone iodine at a final concentration of 0.024% effectively inhibited growth. This is in accordance with findings by Abdolrasouli et al. (2017), who reported MICs between 0.07%-1.25%. Disinfectants containing 0.05% sodium hypochlorite are effective against clade I *C. auris* but not clade IV (Erganis et al., 2014). The growth of *C. auris* UOFS Y-4024 was inhibited by 0.012% sodium hypochlorite solution. Similar to other studies, we found that chlorhexidine combined with isopropyl alcohol is more effective than chlorhexidine as the sole active ingredient (Moore et al., 2017; Rutala et al., 2019; Johnson et al., 2021). A range of quaternary ammonium compounds (QACs) were also tested and found to be very effective at low concentrations. This observed sensitivity to QACs is interesting, as many studies have indicated that *C. auris* is not susceptible to this class of compound (Caceres et al., 2019; Voorn et al., 2023; Erganis et al., 2024). It is known that contact time may influence the effect of different disinfectants (Voorn et al., 2023), making it difficult to compare results between studies. The use of 20 minutes contact time in our study may, therefore, have decreased the reported MICs compared to those reported in literature.

**Table 3.** Minimum inhibitory concentrations (MICs) of commercial disinfectants against *C. auris* UOFS Y-4024.

| Active compound | MIC (% final concentration) |
| --- | --- |
| Ethanol | 29.5 |
| 10% povidine iodine | 0.024 |
| 3.5% sodium hypochlorite | 3.52 |
| 12 % polydimethylammonium chloride | 0.006 |
| 80% didecyldimethylammonium chloride | 0.0012 |
| 50% benzalkonium chloride | 0.003 |
| 5% chlorhexidine gluconate | 3 |
| 0.5% chlorhexidine gluconate combined with 70% isopropyl alcohol | 0.25 |

### Clade assignment of *C. auris* UOFS Y-2024

The whole genome sequence was deposited in NCBI BioProject database (BioProject ID: PRJNA1214514). Whole-genome phylogenetic analysis identified the *C. auris* strain as belonging to clade III (Figure 2).

**Figure 2.**
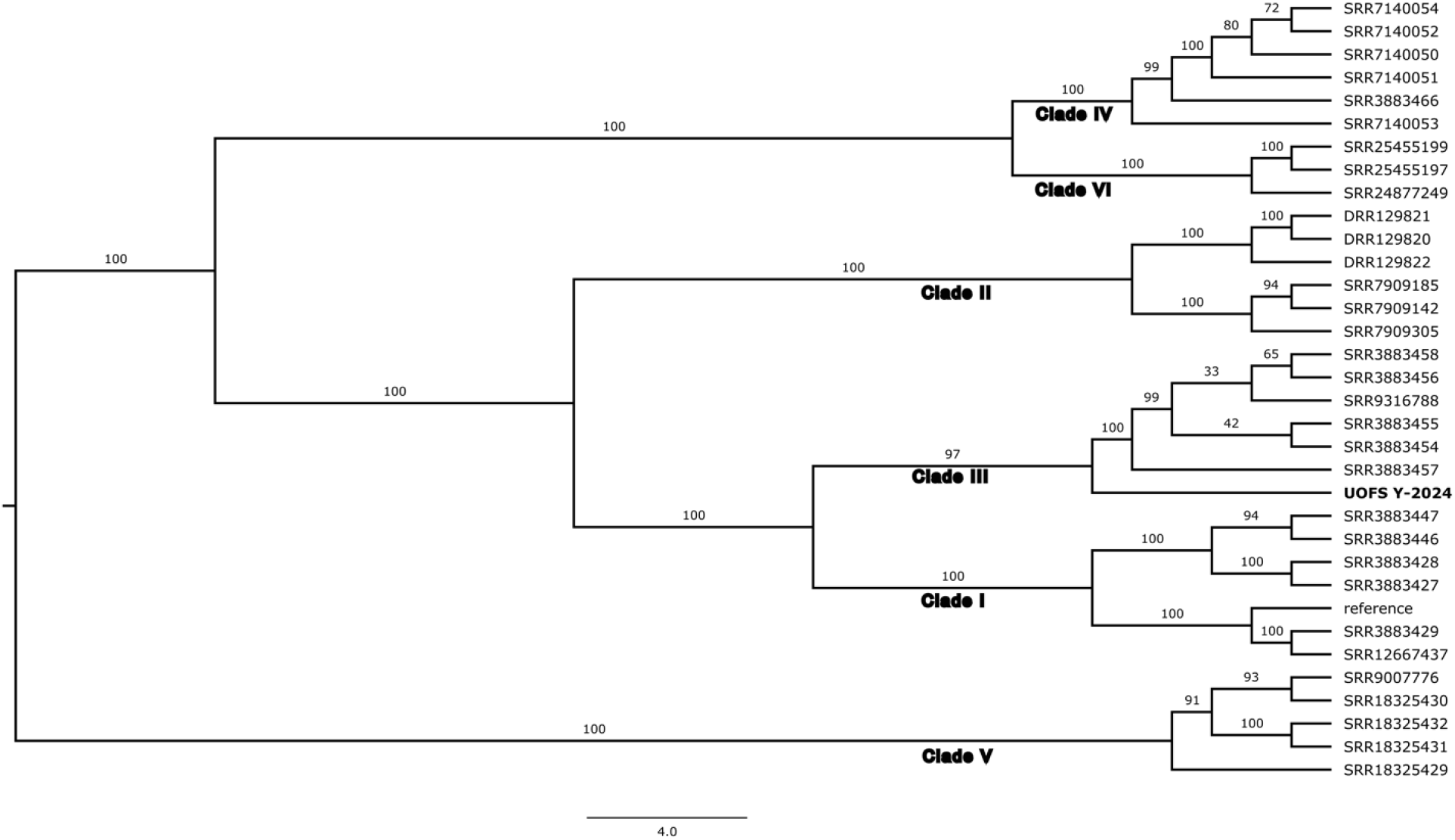
A whole genome SNP-based phylogeny produced using rapid neighbour-joining within the mycoSNP pipeline. The newly isolated *Candida auris* sample (UOFS Y-2024 – in bold) was identified as a member of Clade III using RapidNJ, FastTree and QuickSNP. Samples other than UOFS Y-2024 and the reference sequence (see text) are representative samples from Clades I to VI with identifiers from the Sequence Read Archive. Values on branches represent the percentage of bootstrap support.

## Conclusions

This study highlights for the first time the presence of *C. auris* in the digestive tract of the brown locusts, *L. pardalina*, and shows their potential in disseminating this emerging pathogen. Isolation from the fore- and hindgut implies that the fungus is acquired through feeding and may survive the digestive process and re-enter the environment via faeces. The ability of *C. auris* UOFS Y-4024 under high salt concentration and high temperature correlatess with the environmental and physiological tolerance of locusts, while its resistance to fluconazole and susceptibility to other antifungals are similar to those of clinical and non-clinical strains. The sensitivity to a range of disinfectants, including ethanol and QACs enable practical approaches to containment, although the possibility of acquired resistance to these disinfectants is concenrning. Interestingly, the fact that this strain belongs to Clade III, which is dominant in hospitals in South Africa, raises the questions if thermophilic insects may be a source of clinical strains, or if environmental contamination by clinical strains, could lead to the presence of this yeast in insects. In conclusion, the ability of *C. auris* to survive various environmental and chemical stressors in the digestive system of locusts highlights the importance of understanding the interaction between insects and pathogeninc yeasts and the possible role of insects in the emergence of new pathogenic yeasts under different ecological regimes. This is especially important as locusts (and other insects) are often harvested as food source in many parts of the world (Figure 3).

**Figure 3.**
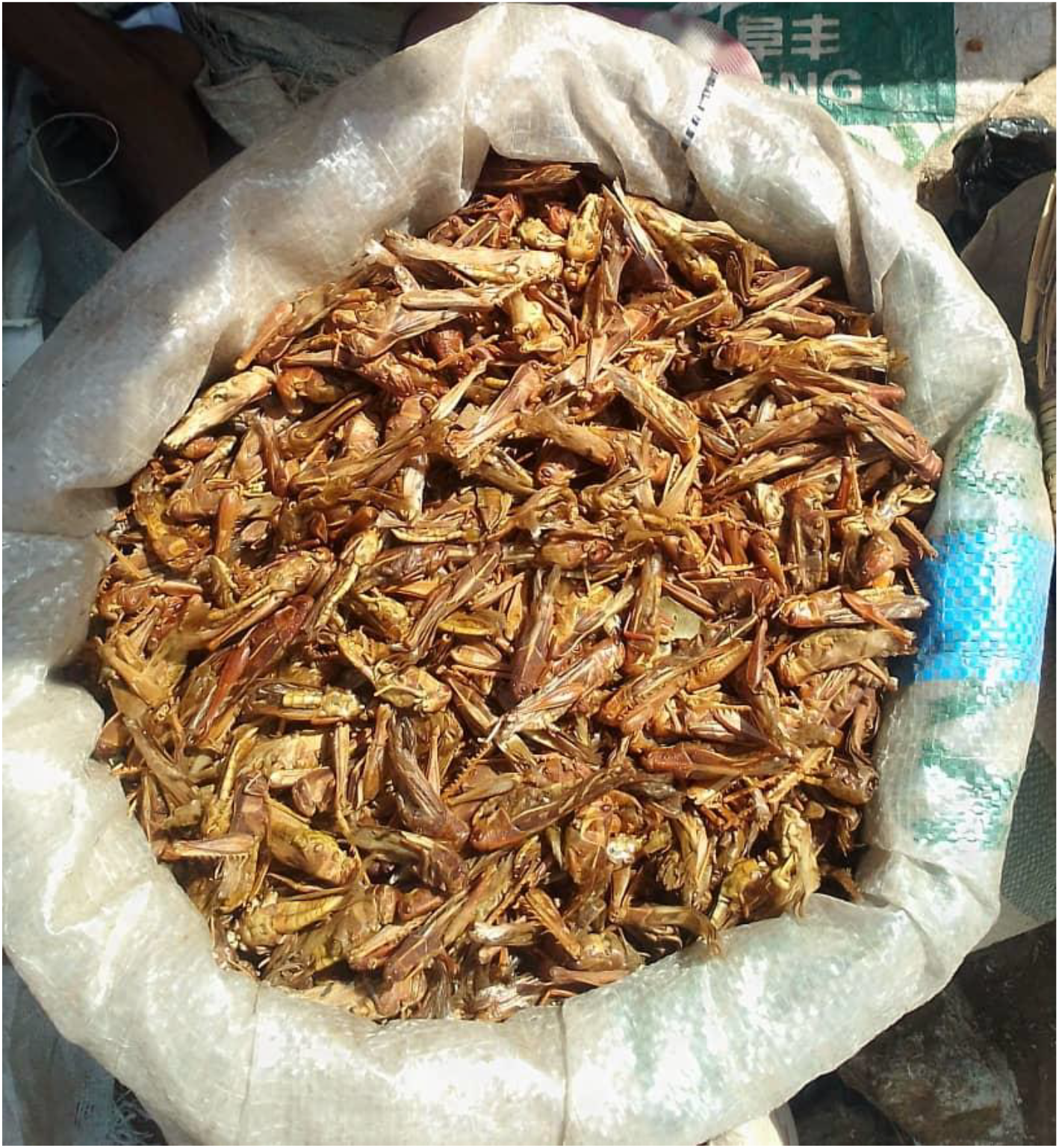
Bag of locusts for sale as food in a market in Nigeria

## Funding

This study was funded by the National Research Foundation of South Africa, under the Foundational Biodiversity Information Small Programme (grant number FBIS2204062196 to CHP).

## Acknowledgement

Tha authors wish to acknowledge Dr Sunday Ayuba Buru for providing the photo for Figure 3.

## Notes

### Competing Interest Statement

The authors have declared no competing interest.

